# Resequencing and association mapping of the generalist pathogen *Botrytis cinerea*

**DOI:** 10.1101/489799

**Authors:** Susanna Atwell, Jason A. Corwin, Nicole Soltis, Wei Zhang, Daniel Copeland, Julie Feusier, Robert Eshbaugh, Daniel J. Kliebenstein

## Abstract

We performed whole genome resequencing of 84 field isolates of *Botrytis cinerea*, largely collected from a local set of plant species. Combined with 13 previously resequenced isolates sampled from diverse locations, this gave a collection of 97 isolates for studies of natural variation. Alignment to the reference sequence T4 and SNP detection provided further data for population genetics analysis including a mapping population for association studies. Although much of the genomic diversity was captured in the original 13 isolates, the additional genomes increased total diversity in the population by a third. Surprisingly, the same additional genomes increase mitochondrial diversity 2-fold. Across the population, LD was limited and decayed rapidly, reflecting frequent outcrossings. Effectively, this sampling strategy increased the level of genetic diversity available, whilst limiting the problem of population stratification and enabling GWAS of several phenotypes on common *Arabidopsis* plants affected in disease pathways. Overlap of results using all GWAS methods revealed numerous candidate genes / pathways that potentially contribute to its broad host range and offer conceivable pathogen decrease targets.

## Introduction

Interactions between plants and pathogens are a key factor in crop yield, with a large body of research studying its evolutionary and mechanistic basis. The majority of plant pathogen research has however focused on specialist pathogens that infect a small number of economically significant species and frequently cause epidemic bursts of disease. This specialization of epidemic pathogens to limited hosts has generated a model of pathogen evolution that is dominated by the generation of new large-effect virulent loci in the pathogen that are countered by equally large-effect resistance alleles in the plant. In contrast to epidemic specialist pathogens are the endemic generalist pathogens such as *B. cinerea* and *S. sclerotorium*. The isolates of these pathogens have the ability to infect nearly any plant host and are widely distributed in the natural environment. However, the evolutionary and genomic pressures that are imparted by this generalist endemic life-style are less well understood.

*Botrytis cinerea* is an endemic necrotrophic plant-pathogenic fungus, the causal agent of gray mold. As a generalist fungus, it infects most vegetable and fruit crops, and a large variety of shrubs, flowers, trees and weeds, including many crops of economic importance (Dean et al. 2012). It is the most important postharvest decay pathogen (Romanazzi and Feliziani 2014) and the second most economically significant plant pathogen (Dean et al. 2012). Whilst there is some evidence for host specialization / increased virulence by some isolates of *B. cinerea* on specific host species (Cotoras and Silva 2005), the effects of geography, host selection and agronomic practices on diversity are poorly understood. Field collected isolates are almost entirely haploid and the presence of a small sequenced genome makes this pathogen a useful model to investigate how a generalist endemic life cycle may structure the genomic variation in the pathogen.

A key component of the evolutionary pressures on generalist pathogens is the interplay of resistance and virulence mechanisms between the pathogens and the host. It is well known on the plant side that resistance to generalist pathogens is typically dominated by a highly polygenic architecture involving 100s to 1000s of genes (Corwin and Kliebenstein 2017; H. C. Rowe and Kliebenstein 2008). To date, there are only a limited number of highly successful quantitative genetic studies of pathogen traits suggesting that virulence can be affected by multiple genes in specialist pathogens (M. H. Lendenmann et al. 2016; Mark H. Lendenmann, Croll, and McDonald 2015; Mark H. Lendenmann et al. 2014). Similarly, there is a wide range of known *B. cinerea* pathogenicity genes / effectors /gene clusters and potential pathways such as Botrydial and Botcinic acid, Velvet, PolyKetide Synthases (PKS) and Non Ribosomal Peptide Synthetases (NRPS) suggesting the potential for polygenic variation in *B. cinerea* virulence (Dalmais et al. 2011; Pinedo et al. 2008; Schumacher et al. 2012). However, we have little understanding if generalist pathogen virulence is equally polygenic to the host plant.

While Genome-wide association studies (GWAS) are one approach that has been particularly useful in uncovering polygenic architectures, especially in species without structured mapping populations, the number of GWAS in necrotrophic pathogens have been few. Additionally, these studies have largely been limited either by low SNP coverage (Gao et al. 2016; Talas et al. 2016) or small genotype sample (Dalman et al. 2013; Palma-Guerrero et al. 2013; Wang et al. 2018). To utilize GWA to understand the architecture of virulence, we developed a large GWAS population within the generalist endemic pathogen *B. cinerea*. This provides a more detailed image of the polygenic genomic architecture of virulence variation.

We performed whole-genome resequencing of 84 isolates of *B. cinerea* to enhance our understanding of variation in this pathogen. This data, combined with 13 previously resequenced isolates from our pilot study, found high levels of polymorphism, including 1,516,518 unique SNPs when mapped against the ~ 41 Mb of the reference sequence from the *B. cinerea* isolate T4. Similar levels were found when the isolate genomes were aligned to an additional *B. cinerea* isolate reference BO5.10. We describe the genomic pattern of polymorphism, the population structure and low levels of linkage disequilibrium (LD) decay among the isolates sampled, loss of function mutations and selective sweeps found. The SNP data additionally enabled the creation of GWAS mapping populations of *B. cinerea* that were phenotyped on 4 genetic backgrounds of *Arabidopsis thaliana*, Col-0, *npr1, coi1* and *pad3*. Using multiple GWA mapping approaches with different SNP sets identified a highly consistent picture of virulence variation in the pathogen. Specifically, virulence is highly polygenic within *B. cinerea*. Further, the identified candidate genes were predominantly dependent on the hosts resistance mechanism(s) but there were some general virulence loci. Thus, the genetic architecture is highly dependent on the interaction of the host and pathogens genomes. The results identified potential targets to assist in future control of this pathogen.

## Mat&Methods

### Polymorphism Detection

DNA was purified from 97 single-spore derived *B. cinerea* isolates as described previously (Atwell et al. 2015). Genomic sequencing, sequence clean-up, alignment of isolates against both the T4 and later the BO5.10 reference genomes and variant calling was carried out as described previously in (Atwell et al. 2015). Analysis was mostly carried out using the T4 data. Additionally, alignment and SNP calling to the BO5.10 mitochondrial and later the genomic reference sequence, that was published during our analyses (JA Kan Van et al. 2017), was also conducted using the same methodologies. However, while individual SNP and indel statistics were determined from separate isolate specific vcfs using SnpEff (Cingolani et al. 2012) for the pilot study, the statistics for these larger datasets were determined using VcfStats v1.2 (Lindenbaum 2015) and rtg-tools-3.8.3 (Cleary et al. 2015) as a group. While SNPeff and VcfStats/rtgtools give the same total number of indels and SNPs for the isolates as a whole, individual indel numbers deviated from the pilot study. As both VcfStats and rtgtools gave accurate results for our lab BO5.10 isolate aligned to the reference sequence for both genomic and mitochondrial sequences and near identical scores to each other, we consider the pilot study statistics for indels an underestimate.

### Phylogenomic analysis

Consensus sequences were created from all reference-based alignments and converted to fasta format using GATK (McKenna et al. 2010). A distance-based approach was used for phylogenetic inference. BIONeighbor-joining radial phylogenetic trees were reconstructed from pair-wise distances using SEAVIEW (Gouy, Guindon, and Gascuel 2010) and plotted using Figtree (Rambaut, A. and Drummond, A 2008). This revealed that 11 genomic aligned isolates sampled from an organic field were highly similar. As such further analysis was conducted using 87 isolates, including a single representative isolate from the 11 highly similar isolates.

### Population structure

We tested for the presence of population structure and admixture using STRUCTURE 2.3.4 (Pritchard, Stephens, and Donnelly 2000) on a filtered SNP subset with a minimum interval of 500bp between SNPs. STRUCTURE was run in the admixture model and 1 to 10 genetic clusters (*K*) were tested, repeated ten times per *K* with a burn-in of 1000 iterations followed by 1000 MCMC iterations. To estimate the most likely value of *K*, the likelihood of the data for a range of *K* values was calculated by creating posterior probabilities of *K*. The different Q matrices were merged with CLUMPP (Jakobsson and Rosenberg 2007) and plotted with strplot (Ramasamy et al. 2014).

### Principle Component Analysis (PCA) of Isolate Variation

PCA was conducted using pca2d (Khang et al. 2016) on the 87 isolates aligned to T4 based on geography (origin of sample location) and family (plant family from which the isolate was sampled).

### Population Genetic Statistics

Haplotype diversity, Nucleotide diversity and ZnS were calculated using PopGenome (Pfeifer et al. 2014) and SNPs from the 87 isolates with a minor allele frequency of at least 0.1 and complete information in nonoverlapping 5-kb windows. Decay in linkage disequilibrium (LD) was determined using PopLDdecay (C. Zhang et al. 2018) with the collection of 87 isolates. Additional subsets of isolates grouped by host plant family or location of isolate were also utilized to assess for differential decay. Selective sweeps were determined from the 87 isolates using SweeD_v3.3.2. (Pavlidis et al. 2013), a likelihood-based detection method of selective sweeps based on allele frequencies which was used to identify candidate genomic regions that are indicative of strong selection pressure at the population level. The composite likelihood ratio (CLR) was calculated for each contig with a resolution of 5kb bins.

### Virulence Phenotyping

The virulence of the collection of *B. cinerea* isolates on the *Arabidopsis thaliana* (*A. thaliana*) genotypes (Col-0, *pad3, npr1, coi1*) was previously measured using a detached leaf assay that allows for high-throughput analysis and is typically consistent with whole plant assays (Boydom and Dawit 2013; Denby, Kumar, and Kliebenstein 2004; Govrin and Levine 2000; Mengiste et al. 2003; Mulema and Denby 2012). The virulence measurements for Col-0, *coi1-1* and *npr1-1* were obtained from the previous publication (W. Zhang et al. 2017). In the same experiments as previously reported for these three genotypes, we had also included the *Arabidopsis* genotype *pad3* that is deficient in camalexin synthesis, a major defense metabolite against *Botrytis*. Pathogen inoculation and utilization of digital images to determine lesion area protocols were as described previously (W. Zhang et al. 2017). Lesion data was analyzed using a generalized linear model with a Gaussian link function for lesion area in an individual model for each plant genotype as described previously (W. Zhang et al. 2017). Using this model, we calculated the least-squared means of the phenotypes within each plant genotype as the phenotype input for GWAS.

### GWAS mapping population

For GWAS mapping, the vcf files of SNPs from the 87 isolates aligned to the reference T4 and separately to the reference BO5.10 and SNPs were filtered (minimum 6x coverage, minimum quality score of 30 and MAF 0.2) and formatted to tab separated files using vcftools (Danecek et al. 2011). This gave 345,485 loci from isolates aligned to the T4 reference and GWAS analysis was conducted using the methods Wilcoxon (Wilcoxon 1945) and bigRR (Shen et al. 2013).

Because bigRR provides an estimated effect size, but not a p-value, we performed permutation analyses to determine effect significance. We permuted the phenotypes 1000x and re-ran the analysis, to establish 95%, 99%, and 99.9% thresholds for significance. GWA analysis was also carried out with GEMMA (X. Zhou and Stephens 2012) using 237,878 SNPs, at MAF 0.20 or greater, aligned to the BO5.10 reference, and less than 10% missing SNP calls as described above. To determine significance of SNPs by GEMMA and maintain consistent methods with the bigRR results, we also used 1000 permutations to determine p-value significance at the 99%, and 99.9% thresholds (Corwin et al. 2016a; Doerge and Churchill 1996; Shen et al. 2013).

From the significant associations, candidate gene lists were created by including genes found as having a candidate SNP within the overlap of the top 500 SNPs in both the Wilcoxon and bigRR mapping methods and were annotated to specific genes using SNPdat (Doran and Creevey 2013) and the T4 gene models for genomic DNA (broadinstitute.org, (Staats and Kan 2012a). The resulting genes were compared to the results from the BO5.10 mapping population GEMMA using annotation from Inra http://dx.doi.org/10.15454/IHYJCX

## Results

### Isolate sampling

To investigate the genomic architecture of genetic variation within the broad host-range generalist *B. cinerea*, we obtained a collection of isolates using single-spore isolation approaches. This collection of natural isolates was obtained mainly from California with a sampling of available international germplasm (Supplemental Table 1). Previous work on a subset of this collection has shown that this represents a population displaying a wide range of genomic, phenotypic and virulence variation (Atwell et al. 2015; Corwin et al. 2016b; Heather C. Rowe and Kliebenstein 2007). By collecting and extending the population using a local set of diverse plant species in combination with a sampling of isolates available from around the world, we worked to minimize extensive linkage disequilibrium (LD) and population structure. This strategy should increase the level of genetic diversity that is available, while limiting the problem of population stratification.

### Genomic diversity

Genomic sequencing and polymorphism detection in 84 new isolates was carried out identical to a previous pilot study (Atwell et al. 2015). In combination with 13 isolates from this pilot study, we generated genomic sequencing data on a total of 97 isolates with a mean of 164X-fold coverage per isolate (32–214X, Supplemental Table1). Genome wide polymorphisms in the nuclear genome were identified by alignment first to the previously used sequenced reference T4 (Staats and Kan 2012b). Additionally, alignment and SNP calling to the BO5.10 reference sequence (JA Kan Van et al. 2017), which was published during our analyses, was also carried out to discern if the reference genome affected the observed level and/or patterns of sequence variation. The BO5.10 reference has the advantage of final chromosome designations rather than contigs.

This analysis identified 1,516,518 unique SNPs against the T4 reference; an additional 536,189 SNPs than were found using the 13 isolates from the pilot study (Atwell et al. 2015). The SNP polymorphisms determined against the T4 reference yielded an average of at least one polymorphism between the isolates every 27 bp across all pairwise combinations, giving an average density of 37 polymorphic SNPs per kb within the population. Single isolates had between 127,779 and 392,802 SNPs per isolate against T4. Similarly, the complete collection of isolates identified 396,638 indel polymorphisms, an increase of 161,474 above the original collection of 13 isolates. There was an average of 46,433 (29,003-67,508) insertions per isolate and 37,682 (22,879-57,620) deletions in comparison to the reference genome. Combining the polymorphisms (both SNP and indel) from all pairwise combinations of isolates against the T4 reference yielded an average of at least one polymorphism between the isolates every 23bp, giving an average density of 43 polymorphisms per kb within the population. There was no significant correlation between polymorphism per isolate and depth of sequencing per isolate (Supplemental Table1).

To test the influence of the reference genome on these estimates, we also mapped the resequencing data against the BO5.10 reference genome (JA Kan Van et al. 2017). This identified a similar rate of polymorphism with 1,673,882 SNPs and 368,981 indels. Single isolates (not including our BO5.10 isolate) had between 165,566 and 402,227 SNPs per isolate, between 14,028 and 65,465 insertions and 12,105 and 56,410 deletions against the BO5.10 reference (Supplemental Table1). This suggests that the two reference genomes give a similar image of a highly polymorphic species. Resequencing the BO5.10 genome using our sample of this genotype identified 125 SNPs against the reference BO5.10 genomic sequence. It is not presently clear if these are mutations or errors in the reference genome or resequencing efforts. Previous work has suggested that spontaneous mutation rates in *B. cinerea* are too low to account for this level of SNPs suggesting that these are technical discrepancies. Given that this rate is >1000 fold lower than the SNP frequency using wild isolates, this indicates that >99.9% of the SNPs identified are real.

### Mitochondrial diversity

Research on quantitative traits in a large number of species has shown the potential for genetic variation in the organelle to influence adaptive traits both alone and in interaction with nuclear polymorphisms (Joseph, Corwin, and Kliebenstein 2015; Paliwal, Fiumera, and Fiumera 2014). To measure variation in the organellar genome, we reconstructed the mitochondrial genome in each of the isolates against the BO5.10 reference mitochondrial sequence. Only the BO5.10 mitochondrial genome was used because the T4 mitochondrial sequence was not published. This found a mean 225X-fold coverage (36–249X) of the 82kb mitochondrial genome and identified 388 SNPs, 1,360 insertions and 669 deletions across the isolates. This is 335 more SNPs than were found in the original collection of only 13 diverse isolates. Single isolates varied between 2 and 288 SNPs per isolate against the reference BO5.10 mitochondrial genome (Supplemental Table 1). Insertion / deletion patterns where similar with individual isolates having between and 5 to 703 insertions, and 1 to 401 deletions (Supplemental Table 1). There was a single SNP and zero indel variation between our BO5.10 sample and the reference sequence (JA Kan Van et al. 2017).

Combining the SNP and indel polymorphisms against the BO5.10 mitochondrial reference yielded an average density of 29 polymorphisms per kb within the population. This showed that mitochondrial genomic variation was approximately 20% lower than for the nuclear genome with an average unique polymorphism every 34bp from the combined isolates. At first impression, this seemed to be a high rate of mitochondrial variation compared to the pilot study (Atwell et al. 2015). However, the resequencing of just 18 isolates of *Lachancea kluyveri* revealed similar levels of polymorphism with an average diversity of 28.5 SNPs/kb and 6.6 indels/kb (Jung et al. 2012). Upon further investigation, it was revealed that this increase in our mitochondrial sequence variation was due to 3 isolates; 1.03.20 with a dramatically higher SNP variation and Peachy and Rasp with greatly increased numbers of indels compared to the other isolates that were sequenced.

A closer look at mitochondrial sequence of the two isolates with increased indel number revealed numerous insertions which ranged from 10-32 bp. BLASTing (Altschul et al. 1997) the majority of these insertions returned matches to Botrytis species but 3 came back as potential matches from other species. However, without further de novo mitochondrial sequencing and sequencing of additional Botrytis mitochondria, we cannot determine if these are highly degenerate mitochondria or horizontally transferred mitochondria from other *Botrytis* species.

### Isolate clustering and admixture

To visualize the genetic relationship amongst these isolates, we plotted their phylogenetic relationships. This utilized the nuclear genome-wide SNP diversity obtained from all 97 isolates in combination with a neighbor-joining phylogenetic tree using pairwise SNP differences to T4 to generate the alignments (Figure 1A). The un-rooted radial phylogeny of the 98 isolates (including the reference T4) showed slight clustering and 11 isolates sampled closely from one organic grape field appeared nearly genetically identical. As such all further aligned analysis utilized only 87 isolates, including just one representative isolate from the collection of nearly identical isolates. This phylogeny was similar if the SNPs were called using theT4 or BO5.10 reference genome (Figures 1A and 1B). In contrast to the nuclear phylogeny, the mitochondrial phylogeny (aligned to BO5.10) was dominated by the 3 isolates showing hypervariable mitochondria. We removed these isolates to allow better visualization of the remaining isolates mitochondrial phylogeny (Figure 1C). Together, these trees suggest that there is admixture amongst the isolates as supported by previous evidence of recombination in the genome (Atwell et al. 2015).

**Figure 1.**
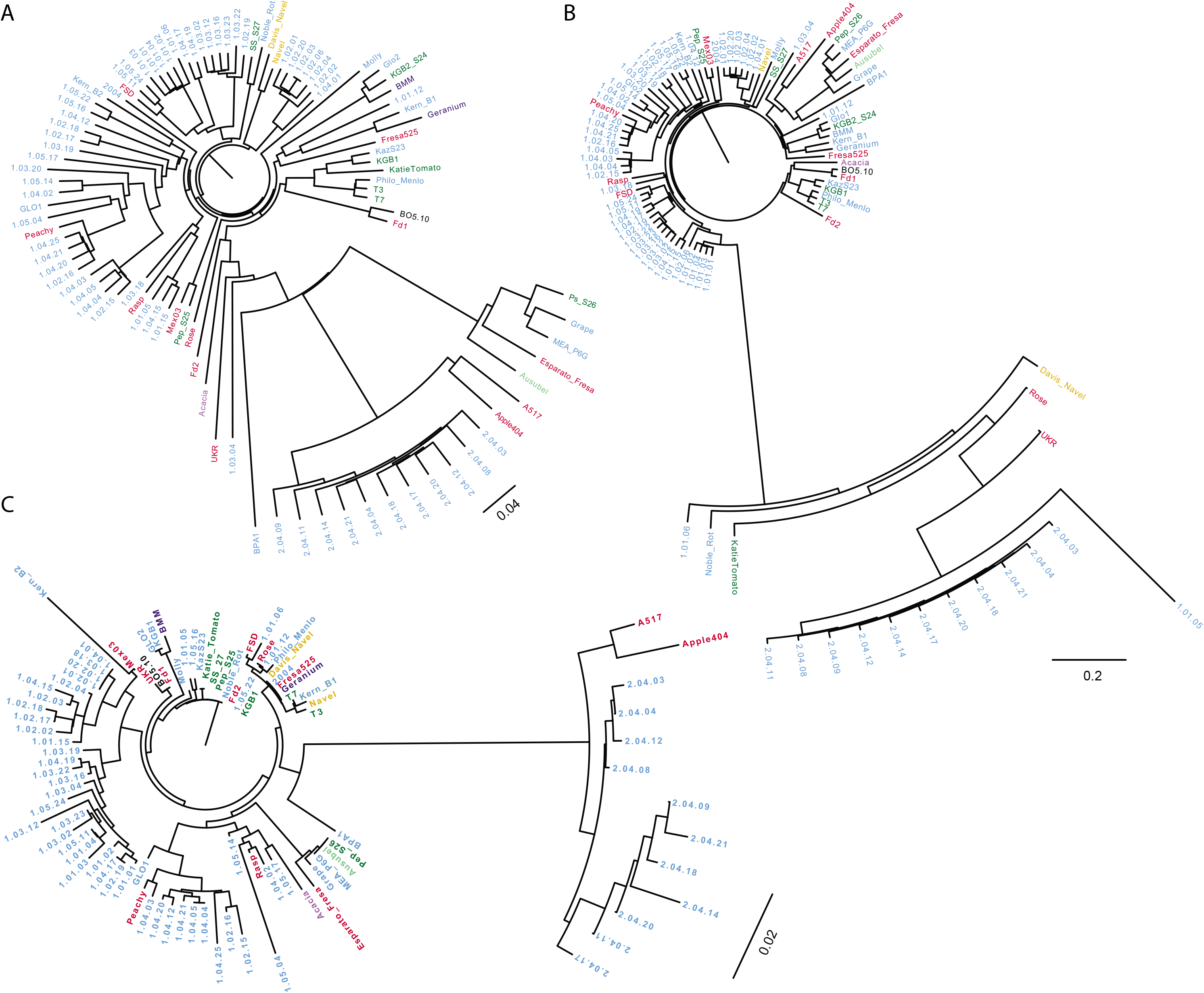
Genomic diversity within 97 isolates of *B. cinerea*. Unrooted genomic phylogeny determined based on pairwise SNP differences in the alignments of **(A)** 97 *B. cinerea* strains to and including the reference T4 and **(B)** 97 *B. cinerea* strains to and including the reference BO5.10. Branch lengths are proportional to the number of segregating sites that differentiate each pair of isolates. **(C)** Unrooted mitochondrial phylogeny determined based on pairwise SNP differences to BO5.10 in the alignments of 97 *B. cinerea* strains (including BO5.10 reference). Isolates Peachy, Rasp and 1.03.20 were removed to enable visualization.

Previous studies with microsatellites and other genetic markers have not clearly resolved the genetic structure of genomic variation in this species. The pilot-study mitochondrial phylogeny suggested the presence of four clusters/populations but the nuclear genomic sequencing of the 13 isolates was less clear. The mitochondrial phylogeny of the larger population (minus the 3 hypervariable isolates) again suggests the presence of four clusters (Figure 1C). Using the nuclear genomic variation with principal component analysis (PCA) while indicating partitioning also indicated some level of admixture with four significant axes that explain ~40% of the genetic variation (Figure 2A). However, none of the vectors show a clear division of isolates by geographical location (Figure 2B) or plant family from which the isolate was sampled (Figure 2C).

To directly assess the presence of population structure and admixture, Bayesian clustering analyses were performed assuming an admixture model using the software STRUCTURE (Pritchard, Stephens, and Donnelly 2000) yielding results consistent with the neighbor-joining phylogeny analyses. From the STRUCTURE and CLUMPP (Jakobsson and Rosenberg 2007; Pritchard, Stephens, and Donnelly 2000) results, *K*=4 (the smallest stable *K* value represents the optimum value) was the best clustering solution of the 87 isolates (Figure 2D). 41 isolates were placed in distinct clusters (made up of 19, 9, 5 and 8 isolates) with the remaining 46 isolates being admixtures of 2-4 of these clusters. This suggests that within our collection of isolates, there is weak population structure that does not appear to map to geography or host plant. Further, half of these isolates represent admixtures of these four theorized populations, which increases the potential utility of genome-wide association mapping.

**Figure 2.**
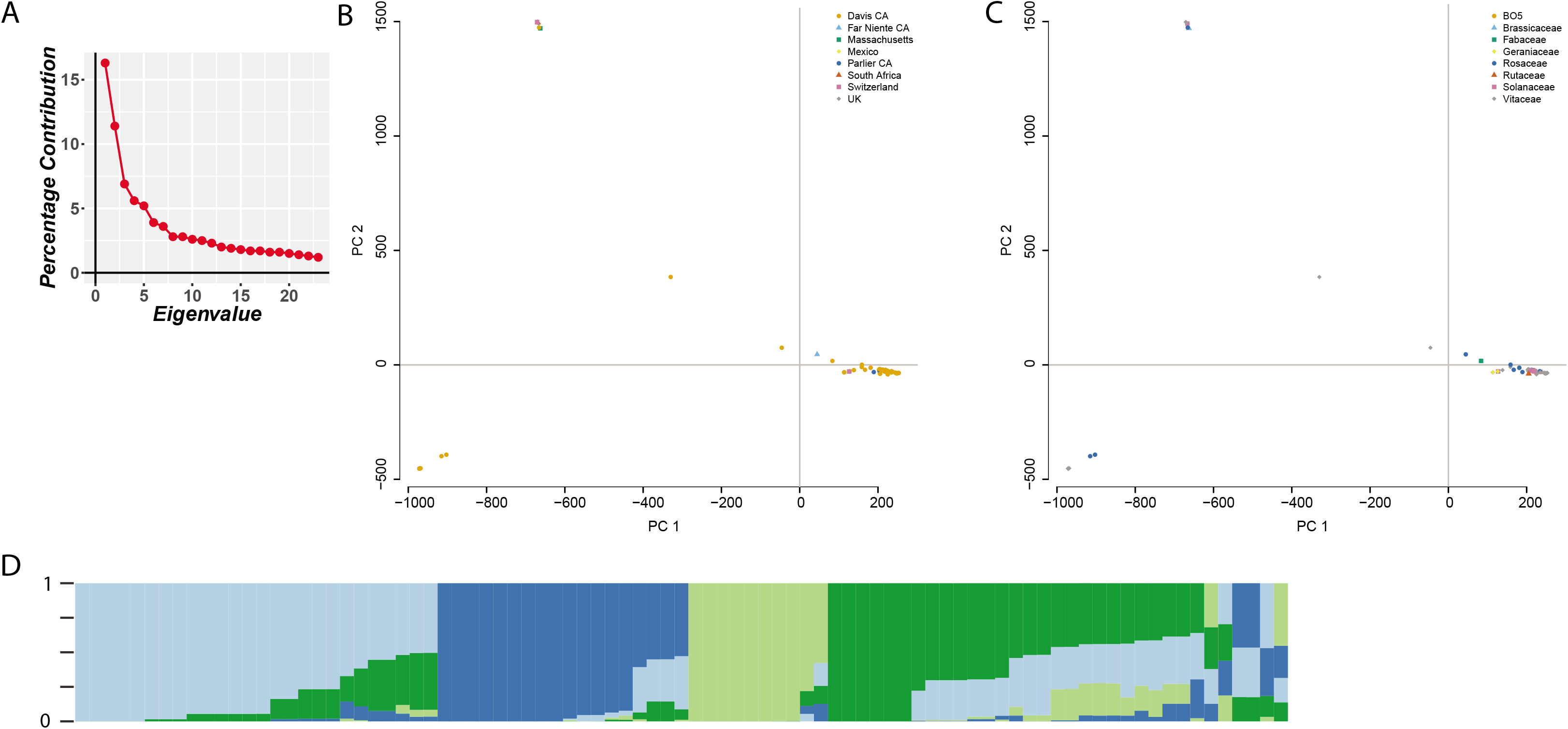
Admixture in *B. cinerea*. **(A) Scree plot** of principal components extracted from all SNP genotypes simultaneously, showing variance contribution based on their eigenvalues. **PCA of genotypes** does not show any clustering by **(B)** geographic or **(C)** plant family origin, suggesting admixture. **(D) strplot** (Ramasamy et al. 2014) showing admixture between 4 ancestral populations from STRUCTURE (Pritchard, Stephens, and Donnelly 2000) results and using *CLUMPP* (Jakobsson and Rosenberg 2007) to align the ten replicates for *K* = 4 (all runs were performed with 1000 burnin period and 1000 MCMC repeats after burnin).

### Linkage Disequilibrium

Structure analysis suggests the presence of extensive admixture among the isolates. This result is supported by previous work with genomic sequencing and AFLPs (Atwell et al. 2015; Walker et al. 2015) that suggested the potential for extensive recombination within this fungal species. To test how this may influence genomic variation within this larger collection of isolates, we measured the decay of linkage disequilibrium across the genome. Across most of the contigs, LD decayed rapidly and mean r2 across the genome fell to approximately one-half of the initial value within 1 kb (Figure 3A). This further supported the presence of relatively frequent outcrossing and recombination within this species. As has been found in a number of other species, mean LD was variable (0.0128 – 0.827, ZnS (Kelly 1997), with some localized regions of high LD (Figure 3B and 3C). A closer look at LD decay by grouping the isolates by the plant family from which they were isolated or the international isolates by themselves showed similar rates of LD decay as within the entire population (Supplemental Figures 1A-D), additionally supporting that neither host source nor geography is driving genomic architecture. Thus, as noted in the pilot study via recombination breakpoint analysis (Atwell et al. 2015), there is extensive recombination within this species on a genome-wide level, limiting mean LD decay to relatively small regions (< 1,000 bp).

**Figure 3.**
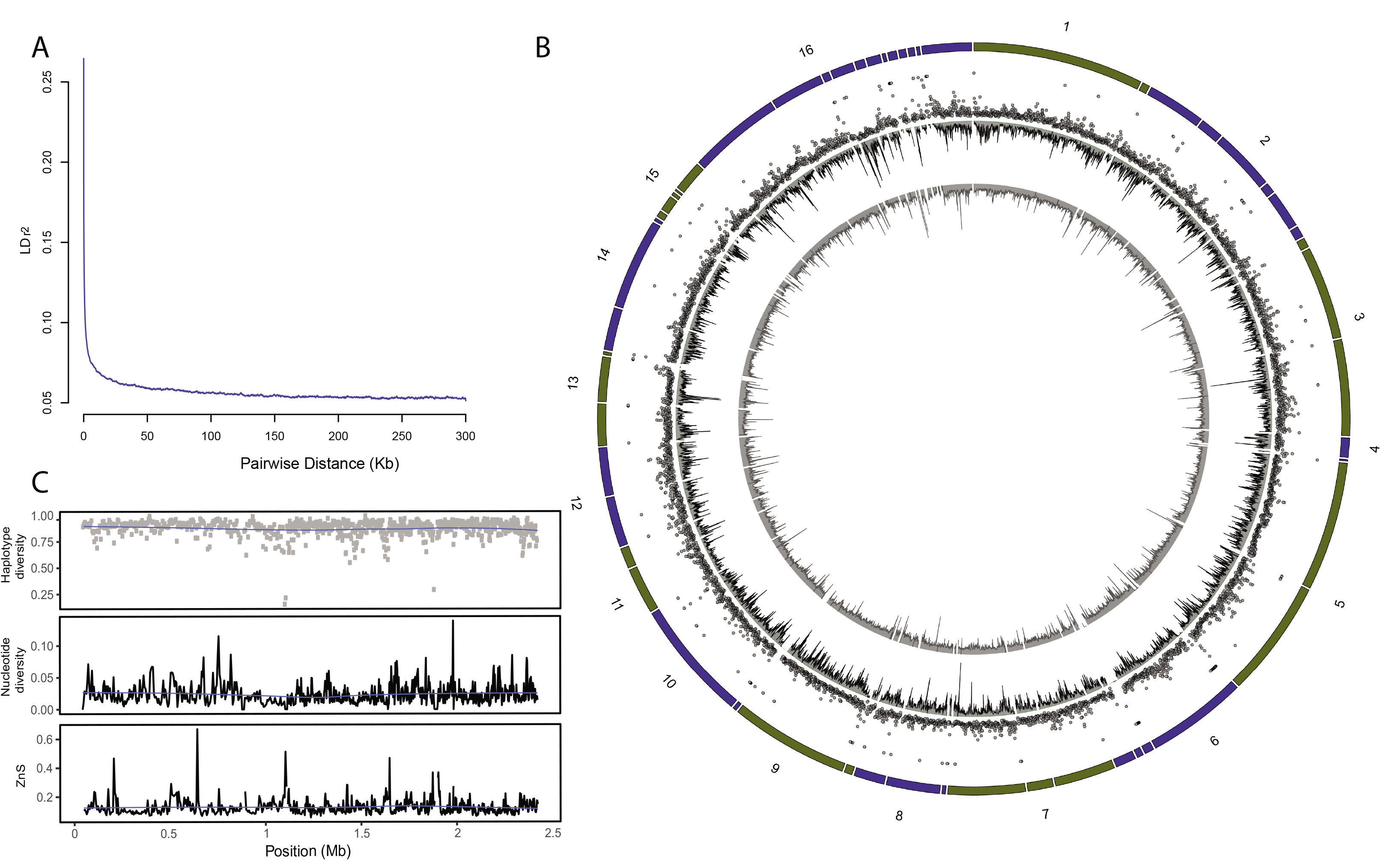
Genome-Wide polymorphism in 87 isolates of *Botrytis cinerea*. **(A)** LD decay was determined by squared correlation of allele frequencies (r^2^) for all pairs of SNPs within the 87 isolates aligned to T4. **(B)** Circos (Krzywinski et al. 2009) diagram of 56 contigs detailing (from outer to inner track) haplotype diversity, nucleotide diversity (Nei, 1987) and mean linkage disequilibrium (Kelly, 1997). Coloring and numbers on the outside of the circle denote potential contig placement on chromosomes for visualization purposes only. **(C)** Close-up of haplotype diversity, nucleotide diversity and mean linkage disequilibrium for contig 2.

### Diversity and Selective Sweeps

The localized regions of higher LD suggested the potential presence of localized variation in selective pressures. To investigate the potential for local variation in sequence diversity, we shifted from a whole genome analysis to discrete local analysis. Across the entire genome, the levels of nucleotide diversity estimated in non-overlapping 5-kb windows ranged between 0.0001 and 0.044 per site amongst the 84 isolates (Figure 3B and 3C), with a mean nucleotide diversity of 0.005 across all combined contigs. Measurements of haplotype diversity in nonoverlapping 5kb blocks gave elevated but not unusual levels ranging between 0.02 and 0.99, with an average of 0.85 (Figure 3B and 3C). Thus, there is extensive local variation around the mean genomic sequence diversity.

One potential process that can alter local sequence variation are selective sweeps that are indicative of strong directional selection pressure at the population level. The presence of low levels of genome-wide haplotype diversity warranted investigation of selective sweeps. Analyzing the genomic variation with SweeD (Pavlidis et al. 2013), identified 56 peaks genome-wide with a CLR under the 0.95 criterion calculated with a resolution of 5kbp bins across the whole genome (Supplemental Figure 2).

Similar to the polymorphisms that were predicted to cause major loss-of-function in the pilot study, these 56 regions identified genes associated with mating and virulence. Particularly heterokaryon incompatibility genes and NACHT domain protein encoding genes that are associated with vegetative incompatibility in fungi (Glass, Jacobson, and Shiu 2000; Glass and Kaneko 2003). Known virulence genes including VELVET, in which the selective sweep is against a minor allele linked to altered virulence and growth (Schumacher et al. 2012), were also found. In addition, there are sweeps affecting Chitin Synthase, Tannase (a lignin degrading enzyme), Patatin and unknown Cytochromes p450. Thus, there appear to be potential selective sweeps in Botrytis that are linked to known mating and virulence loci.

### Genomic and mitochondrial loss of function mutations

To investigate the potential functional consequences of this high polymorphism rate, we extracted polymorphisms predicted to have major effects on gene function using SnpEff (Cingolani et al. 2012) and the annotation from T4 (http://www.broadinstitute.org<u>; *B.cinerea* T4</u>). These could either be via premature protein truncation, triggering of nonsense mediated decay or other major effect polymorphsims. This analysis identified 14,498 polymorphisms likely to have major loss-of-function effects on 2,697 genes (out of a possible 10,401 genes annotated in the T4 reference). This represented an increase of only about 2-fold over the original 13 isolates when adding 74 additional isolates. This suggests that we are beginning to resample the major functional polymorphisms present within the species, but the extensive recombination is allowing us to sample new combinations of alleles.

An analysis of these polymorphisms showed that the vast majority were not singleton polymorphisms as might be expected if they were highly deleterious to the organism. However, half of the loss-of-function polymorphisms had a frequency lower than 20% (Supplemental Figure 3). Importantly, we were able to find the previously reported loss-of-function polymorphism for the VELVET gene (Schumacher et al. 2012) segregating within the population. Additionally, like the selective sweep analysis in this study and the loss-of-function polymorphisms from the previous pilot-study, there was a tendency for loss-of-function polymorphisms to occur in heterokaryon incompatibility genes and NACHT domain protein encoding genes, Tannase, Patatin and Cellulase genes. These control the ability of the isolates to have vegetative mating or to degrade plant cell walls. Thus, there is a large pool of large-effect polymorphisms segregating within Botrytis potentially contributing to its broad host range.

### GWAS and Polygenic Virulence

The linkage disequilibrium and polymorphism data suggest that it should be possible to conduct a GWA study in this population to identify naturally variable *B. cinerea* genes that influence differential virulence on specific hosts. To test this possibility, we individually inoculated all of the resequenced isolates on the common lab *A. thaliana* accession Col-0 and on three *A. thaliana* plants compromised in specific resistance pathways *coi1, pad3* and *npr1*. We obtained average virulence for each pathogen isolate by host genotype combination using eight-fold independent randomized replication across two experiments. The traits followed a normal distribution (Figures 4A-D) except that lesion scoring on Col-0 was slightly right-skewed due to the isolate A517 (Figure 4A). Heritability of virulence across the *B. cinerea* isolates, among the *Arabidopsis* genotypes and interaction between isolate and host genotype have been previously published (H^2^Isolate = 0.164, H^2^HostGenotype = 0.151, H^2^Isolate x HostGenotype = 0.047 respectively; (W. Zhang et al. 2017).

For the GWAS, we utilized up to 345,485 high quality SNPs that were evenly distributed across the *B. cinerea* contigs from the T4 reference and three methods. The first was a Wilcoxon non-parametric SNP by SNP analysis (Wilcoxon 1945). The second was a ridge-regression analysis, bigRR (Shen et al. 2013) that tests all of the SNPs in a single model. This approach has previously been shown to have a high validation rate within *A. thaliana* and the rank order of the SNPs is highly similar to that obtained using the EMMA methodology (Kang et al. 2008). The benefit of the ridge regression is that it can be conducted quickly enough to do direct permutation analysis to find empirical thresholds based on the phenotypic distribution (Shen et al. 2013). Finally, we used a mixed model to control for population structure within the GEMMA (X. Zhou and Stephens 2012) pipeline and a SNP mapping population created against the BO5.10 reference. Using the Wilcoxon and bigRR methods, we identified the genes linked to the top 500 SNPs for virulence on each host genotype (Figure 5, Table 1). We chose the top 500 SNPs, as permutation with bigRR showed that all of these were significant for all traits and we wanted to compare similar sized gene lists across the approaches. SNPs of interest were distributed over all contigs and between 40 and 60 genes were found per phenotype that overlapped between the three methods (Figure 5 and Supplemental Figure 4). Including the GEMMA results showed that the three methods and two mapping populations largely found the same genes or when there was a discrepancy, they identified neighboring genes (Table 1). Using the combination identified between 5 and 9 genes per phenotype as being of immediate interest but these were entirely uncharacterized genes (Table 1).

The *B. cinerea* genes identified as linked to variation in virulence on *Arabidopsis* were highly dependent on the specific host genotype with Col-0, *npr1-1* and *coi1-1* illuminating different pathogen loci. In contrast, *pad3* and *coi1-1* identified largely overlapping sets of virulence associated genes in *B. cinerea*. This similarity between *coi1-1* and *pad3* is possibly because both mutants lead to a loss of production of the major defense metabolite, camalexin, upon infection by these isolates (Schuhegger et al., 2006; N. Zhou, Tootle, & Glazebrook, 1999 (Schuhegger et al. 2006; N. Zhou, Tootle, and Glazebrook 1999). This suggests that the collection of virulence loci found when using the Col-0 genotype but not the *coi1-1* and *pad3* hosts may be associated with how *B. cinerea* counteracts this specific metabolic defense. Additionally, 3 genes were found that are linked to altered virulence on 3 or more host genotypes; NMT1/THI5 like-Thiamin biosynthesis -vit B1 (NMT1), for fungi, thiamine is essential for vegetative growth, development, sporulation, host invasion, and adaptation to environmental stress (Hohmann and A Meacock 1998); Berberine bridge enzyme like (BBE), which is part of the PolyKetide Synthase19 (PKS19) cluster; and a NACHT domain gene. Together, this shows that it is possible to use GWA within the pathogen to find loci that control both general virulence and host-specific virulence mechanisms.

**Figure 4.**
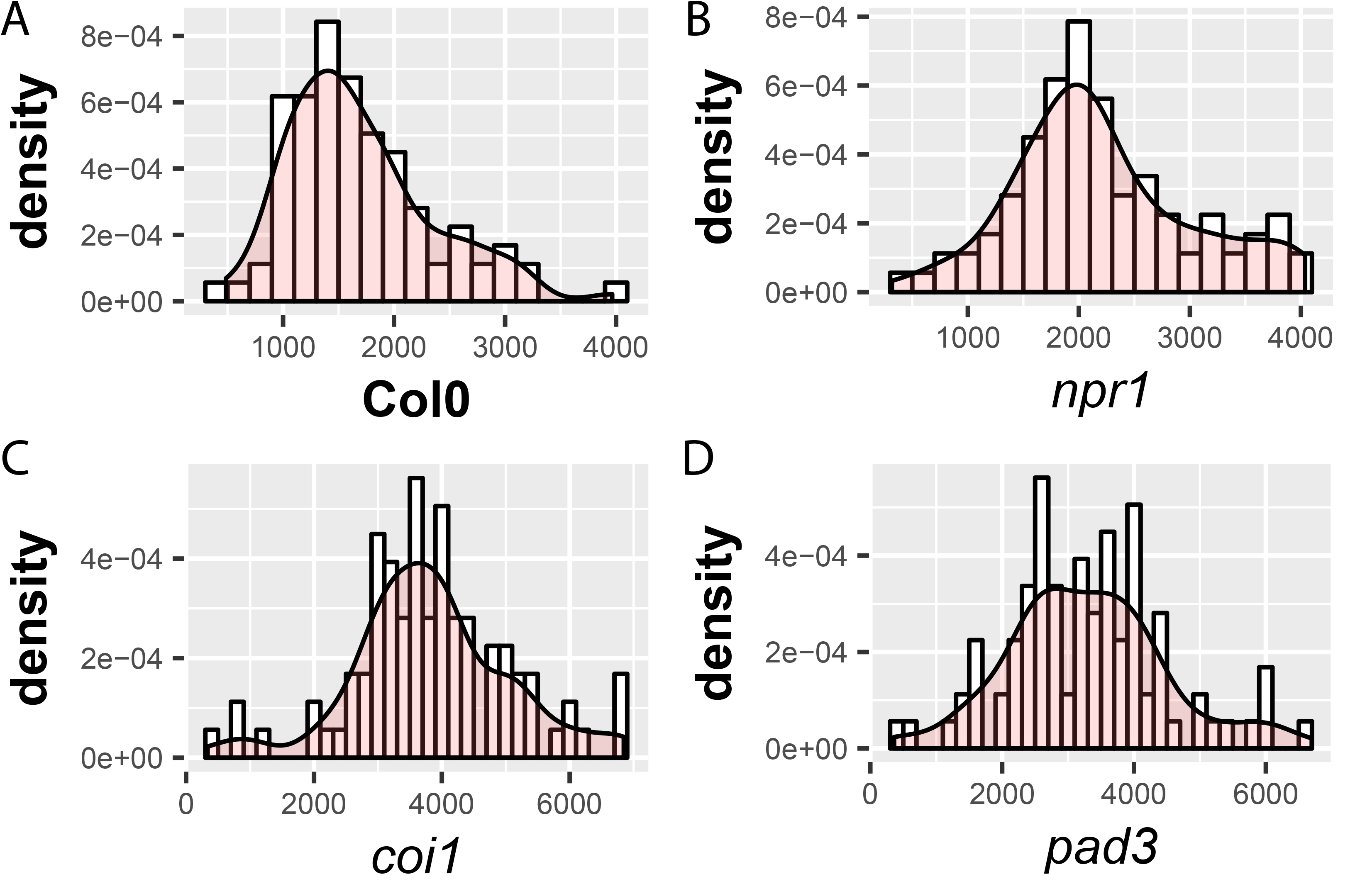
Virulence diversity of the *B. cinerea* isolates on *Arabidopsis thaliana*. Kernal density estimate plots of phenotype distribution. Phenotypes are least-squared means of the lesion area determined following inoculation of each of the resequenced isolates scored on the **(A)** Col-0, **(B)** *npr1*, **(C)** *coi1* and **(D)** *pad3* plant genotypes.

**Figure 5.**
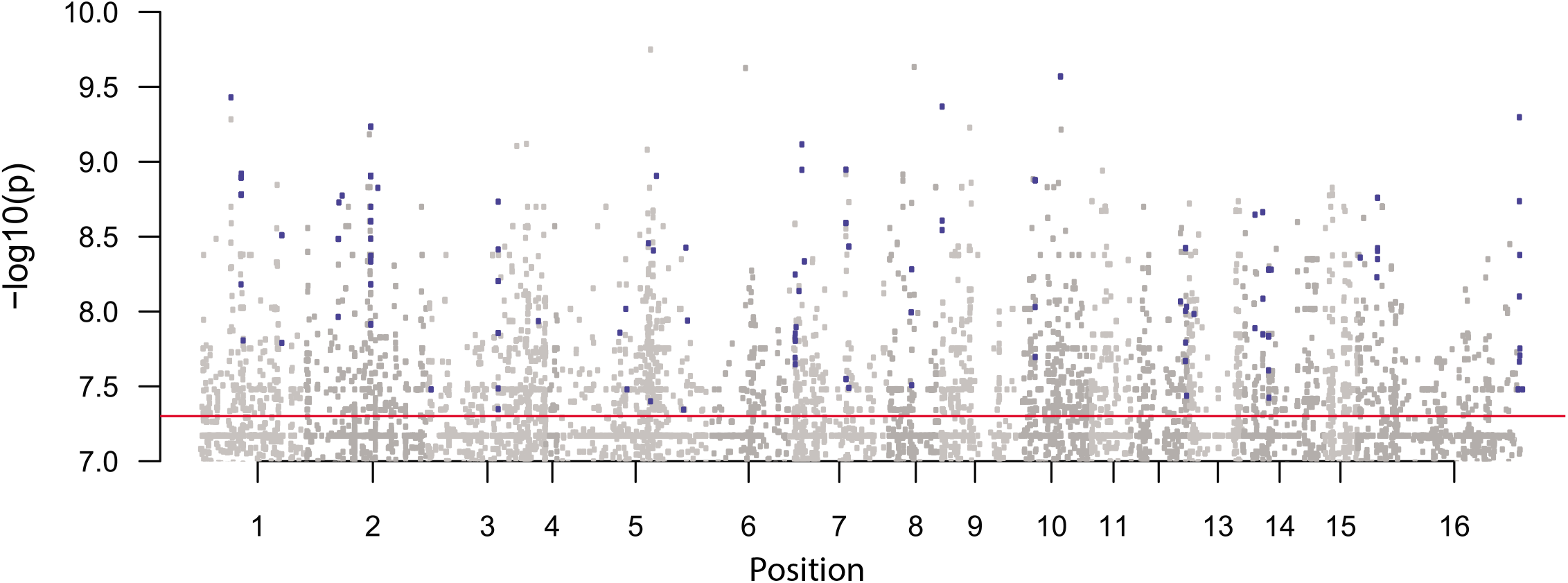
Polygenic architecture of virulence. GWAS Manhattan plot of the Wilcoxon SNP *p*- values from visible lesion phenotypes scored on the Col-0 *Arabidopsis* plant genotype. Negative log10 *p*-values are plotted against genomic position, the blue highlighted SNPs are genes (determined from available annotation) found in the overlap of the top 500 SNPs using all GWAS methods (Wilcoxon and bigRR with SNPs from the T4 alignment and GEMMA with SNPs from the BO5.10 alignment), the red horizontal line denotes genome-wide significance.

**Table 1.** Candidate virulence genes for *B. cinerea* on *A. thaliana*. Genes of interest (determined from available annotation) found in the overlap of the top 500 SNPs using all GWAS methods, Wilcoxon and bigRR with SNPs from the T4 alignment and GEMMA with SNPs from the BO5.10 alignment. Genes are listed by plant genotype on which they were found to be significant.

| Phenotype | Genes of interest found | Annotation | Function / proposed function | BO5.10 annotation | BO5.10 alignment GEMMA overlap |
| --- | --- | --- | --- | --- | --- |
| Col-0 lesion | BcT4_672 | Catalase | protect against host produced active oxgen species /also close to PKS17 | Bcin03g01920 | Annotated and found with all 3 methods |
|  | BcT4_5465 | p450 | Cytochrome P450 |  | Not annotated, next gene BcT4_5464 found |
|  | BcT4_6124 | Rhodanese | Rhodanese-like domain - cyanide degradation to protect against cyanogenic glycosides | Bcin01g00880 | Annotated next genes BcT4_6123 and BcT4_6125 found |
|  | BcT4_6600 | Kelch_4 | Galactose oxidase. CAZy. | Bcin13g05710 | Annotated and found with all 3 methods |
|  | BcT4_10439 | BBE | Berberine and berberine like – part of PKS19 cluster | Bcin16g00006 | Annotated, closest gene found with GEMMA BcT4_10442 |
| npr1 lesion | BcT4_1413 | p450 | Cytochrome P450 | Bcin06g00650 | Annotated and found with all 3 methods |
|  | BcT4_1780 | HET | Heterokaryon incompatibility protein (HET) | Bcin09g03510 | Annotated gene BcT4_1776 found with GEMMA |
|  | BcT4_5465 | p450 | Cytochrome P450 |  | Not annotated, next gene BcT4_5464 found with GEMMA |
|  | BcT4_5925 | Cellulase | Cellulase (glycosyl hydrolase family 5) | Bcin14g01690 | Annotated and found with all 3 methods |
|  | BcT4_7137 | NMT1 | NMT1/THI5 like- Thiamin biosynthesis -vit B1 | Bcin16g00010 | Annotated and found with all 3 methods |
|  | BcT4_9080 | PKS12 | PKS12 | Bcin02g08770 | Annotated and found with all 3 methods |
|  | BcT4_9386 | NACHT | NACHT domain | Bcin04g01020 | Annotated and found with all 3 methods |
| pad3 lesion | BcT4_2611 | EthD | Part of PKS7 cluster | Bcin10g00030 | Annotated gene BcT4_2616 found with GEMMA |
|  | BcT4_3023 | NACHT | NACHT domain | Bcin10g04480 | Annotated and found with all 3 methods |
|  | BcT4_5487 | Fig1 | Ca2+ regulator and membrane fusion protein-fruiting body and ascus development | Bcin15g00760 | Annotated, closest gene found with GEMMA is BcT4_5473 |
|  | BcT4_7137 | NMT1 | NMT1/THI5 like- Thiamin biosynthesis -vit B1 | Bcin16g00010 | Annotated, next gene found with GEMMA |
|  | BcT4_9026 | PKS21 | PKS21 |  | Not annotated, region removed by GEMMA |
|  | BcT4_9386 | NACHT | NACHT domain | Bcin04g01020 | Annotated and found with all 3 methods |
|  | BcT4_9413 | NACHT | NACHT domain | Bcin04g00920 | Annotated and found with all 3 methods |
|  | BcT4_10345 | PKS19 GST_C_2 | Glutathione S-transferase | Bcin08g00300 | Annotated, gene BcT4_10342 found with GEMMA |
|  | BcT4_10439 | BBE | Berberine and berberine like – part of PKS19 cluster | Bcin16g00006 | Annotated, next gene BcT4_10440 found with GEMMA |
| coi1 lesion | BcT4_2611 | EthD | Part of PKS7 cluster | Bcin10g00030 | Annotated gene BcT4_2616 found with GEMMA |
|  | BcT4_3023 | NACHT | NACHT domain | Bcin10g04480 | Annotated and found with all 3 methods |
|  | BcT4_7137 | NMT1 | NMT1/THI5 like- Thiamin biosynthesis -vit B1 | Bcin16g00010 | Annotated and found with all 3 methods |
|  | BcT4_7277 | CAP59_mtransfer | Cryptococcal mannosyltransferase 1- virulence associated enzyme of Cryptococcus | Bcin16g01560 | Annotated, region removed by GEMMA |
|  | BcT4_7646 | NRPS3 Epimerase | NAD dependent epimerase/dehydratase family | Bcin02g02380 | Annotated, closest gene found with GEMMA BcT4_7661 |
|  | BcT4_9026 | PKS21 | PKS21 |  | Not annotated, gene BcT4_9025 found with GEMMA |
|  | BcT4_9386 | NACHT | NACHT domain | Bcin04g01020 | Annotated, closest gene found with GEMMA BcT4_9393 |
|  | BcT4_10056 | HET | Heterokaryon incompatibility protein (HET) | Bcin06g07040 | Annotated and found with all 3 methods |
|  | BcT4_10439 | BBE | Berberine and berberine like – part of PKS19 cluster | Bcin16g00006 | Annotated, closest gene found with GEMMA BcT4_10442 |

## Discussion

Given the economic cost of crop damage caused by *B. cinerea*, this plant pathogen has undergone extensive investigation, with great advancements in predicted gene functionality made recently via sequencing projects (Amselem et al. 2011; Blanco-Ulate et al. 2013; JA Kan Van et al. 2017; Staats and Kan 2012b). One limitation of these studies is that *B. cinerea* is a broad host pathogen and it is unclear as to how much of the species variation, genetic and phenotypic, can be captured solely by two reference genomes and one predominant lab isolate. To address these questions, we conducted whole genome sequencing on a collection of *B. cinerea* isolates, to begin understanding the genomic architecture of this broad host pathogen and how this variation may contribute to this host range. In this study we provide the first comprehensive resequencing data of a collection of *B. cinerea* isolates which enabled the creation of the first *B. cinerea* GWAS mapping population. The availability of this data will be a valuable resource for further *B. cinerea* – plant interactions, identification of genes responsible for traits and potential targets for *B. cinerea* control.

The genomic resequencing identified a high level of genetic variation amongst these isolates including major effect polymorphisms that create natural knockouts in nearly 25% of the annotated genes. This genomic resequencing showed that there was no clear signal of host source or geography as has been found in other studies using AFLPs or microsatellites (Ma and Michailides 2005). Comparing polymorphism identified with the 84 new isolates vs an original set of 13 isolates showed that 2/3rds of the polymorphisms were found in the original 13 resequenced isolates. This suggests that there is a large body of moderate frequency polymorphisms within the species. In combination with the lack of population structure and the rapid LD decay, the new isolates are sampling new allelic combinations more than new genetic diversity as would be essential to maximize the power of any GWAS.

### LD, Life-History and GWA

Fungal pathogens were considered to have a largely clonal life-style similar to Oomycete pathogens (Taylor et al. 2015). This would limit the utility of GWA to identify causal variants influencing variation in virulence across a species. However, recent genomic studies are beginning to show that this is not an accurate view. Recent work has shown that in *S. cerevisiae*, global LD decays within 500bp (Peter et al. 2018), similar to the 1kb found in our study of *B. cinerea*. Smaller, more focused studies in plant pathogens have shown that LD is similar in *Fusarium graminearum* (mean LD ~1 kb, (Talas and McDonald 2015) but higher in *Parastagonospora nodorum* (mean LD ~5-10 kb, (Gao et al. 2016). This could signify that similar pathogens may be more suitable for GWAS than the previous ideas of clonality signified. Our data shows that *B. cinerea* has the combination of high frequency SNPs and low LD required to enable effective GWAS. Further, *B. cinerea* has a lack of substantial sub-structuring suggesting that structure will not be a dominant driver in any GWA study. The fact that the LD decay remained rapid upon dissection of the isolates based on geography or host plant, bodes well for addition of further samples to increase power. This allows for fine mapping of genes which can be targeted to suppress the destruction of this pathogen.

Using this LD and diversity, we tested the potential to use GWA by measuring virulence of the isolates on four defined *A. thaliana* genotypes that differ in their defense capacities. This showed the potential to identify both general and host-specific loci that could influence virulence within this *B. cinerea* population. Using three different GWA approaches within a specific host plant identified largely overlapping sets of potential causal genes regardless of if the method controlled for population structure (GEMMA), tested SNP by SNP (Wilcoxon), or conducted a genome prediction approach (bigRR). Whilst there was overlap in the genes found between the three approaches, the SNPs found within the putative causal genes were not always identical (Table 1). Thus, *B. cinerea* GWA using this population will be an excellent tool to generate gene level hypotheses to follow up.

### GWA across the host genotypes showed some consistency and some host specific genes

Intriguingly, while the results of our GWAS highlighted several genes and pathways of interest we did not find many of the known classical virulence genes. Selective sweep analysis and the identification of major effect polymorphisms showed this is at least partly due to allele frequencies in these classical virulence genes, some of which are below what was included in the SNP mapping populations (<20%). GWAS is usually performed with SNPs, because they have the highest frequency and SNPs of minor frequency can lead to spurious associations (Turner et al. 2011). As such, this suggests that the best way to utilize this population to understand the architecture of virulence and pathogen adaptation is to combine the GWA with a separate analysis that investigates regions like selective sweeps. Together, this combined list of polymorphisms can be used to better develop hypotheses on how a generalist pathogen evolves to a broad array of hosts.

## Supporting information

**Supplemental Table 1. Genetic polymorphism summary**. A summary of genomic sequencing on each isolate. The mean sequencing depth and number of SNPs, insertions and deletions in respect to the T4 and B05.10 genomes for the nuclear genome are listed. The polymorphisms in comparison the BO5.10 mitochondrial genome are also provided. The host and location of collection for each isolate is listed.

**Supplemental Figure 1. Host collection does not influence LD decay**. LD decay was determined by squared correlation of allele frequencies (r^2^) for all pairs of SNPs within each group of isolates sampled from **(A)** Vitaceae, **(B)** Rosaceae, **(C)** Solanaceae and **(D)** international collection sites.

**Supplemental Figure 2. Potential selective sweeps**. Selective sweeps were determined from the 87 isolates aligned to reference T4 using SweeD_v3.3.2.(Pavlidis et al. 2013) which implements a composite likelihood ratio (*CLR*) test. Blue colored SNPs represent SNPs that cross the 0.95 criterion with d=5kbp.

**Supplemental Figure 3. Frequency of major loss of function mutations** from the T4 aligned data.

**Supplemental Figure 4. Polygenic architecture of virulence**. GWAS Manhattan plot of the Wilcoxon SNP p-values from visible lesion phenotypes scored on the **(A)** *npr1*, **(B)** *pad3* and **(C)** *coi1* plant genotypes. Negative log10 *p*-values are plotted against genomic position, the red and blue lines are suggestive and genome-wide significance lines respectively.

