## Supplementary figures and images for "Resequencing and association mapping of the generalist pathogen *Botrytis cinerea*"

### Supplementary file 1

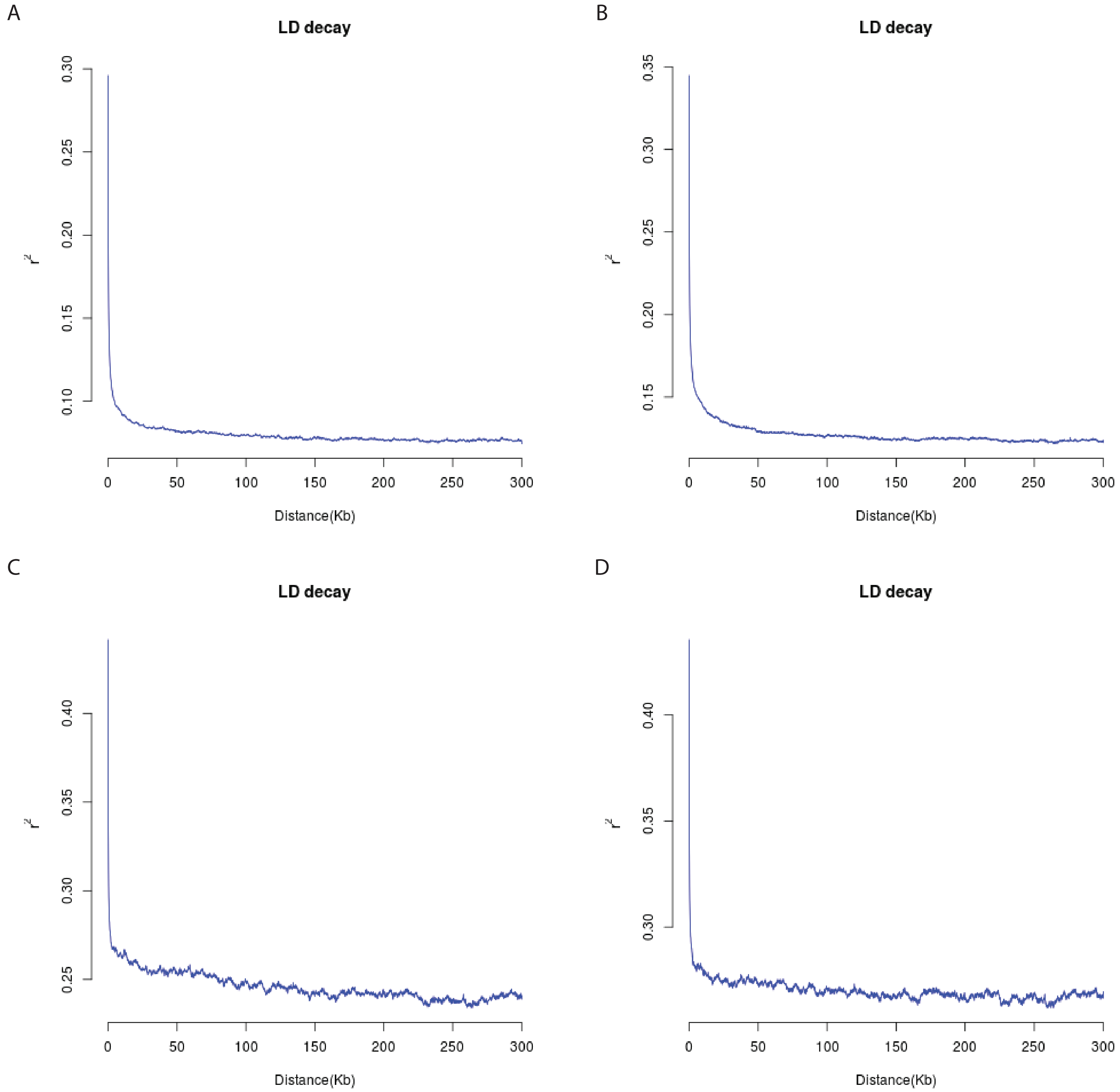

### Supplementary file 2

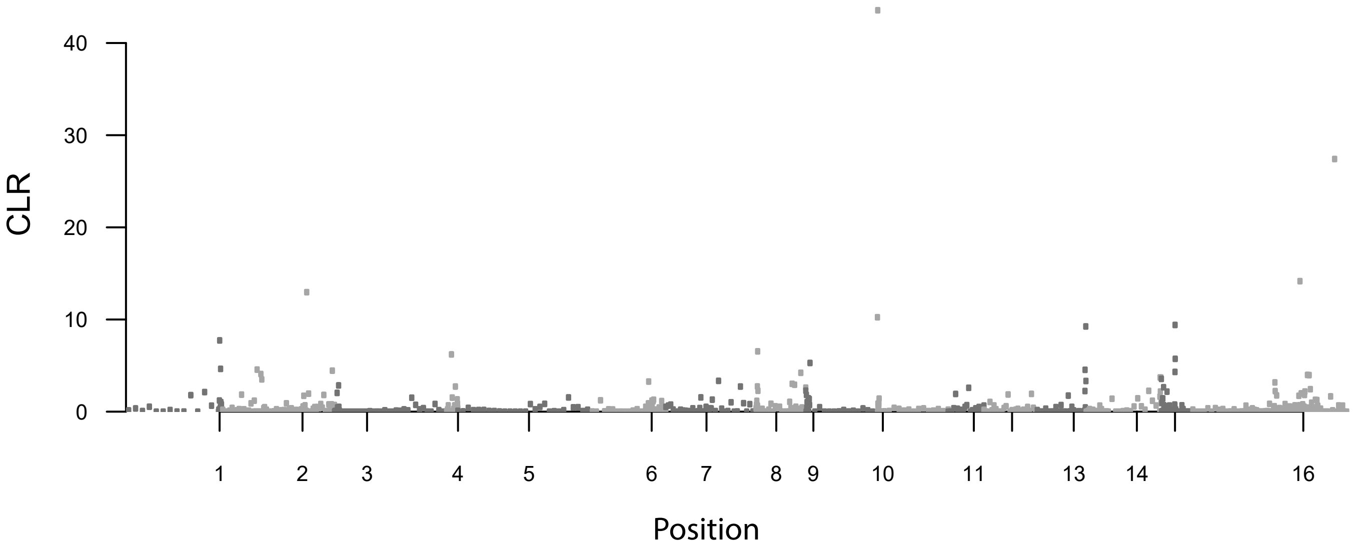

### Supplementary file 3

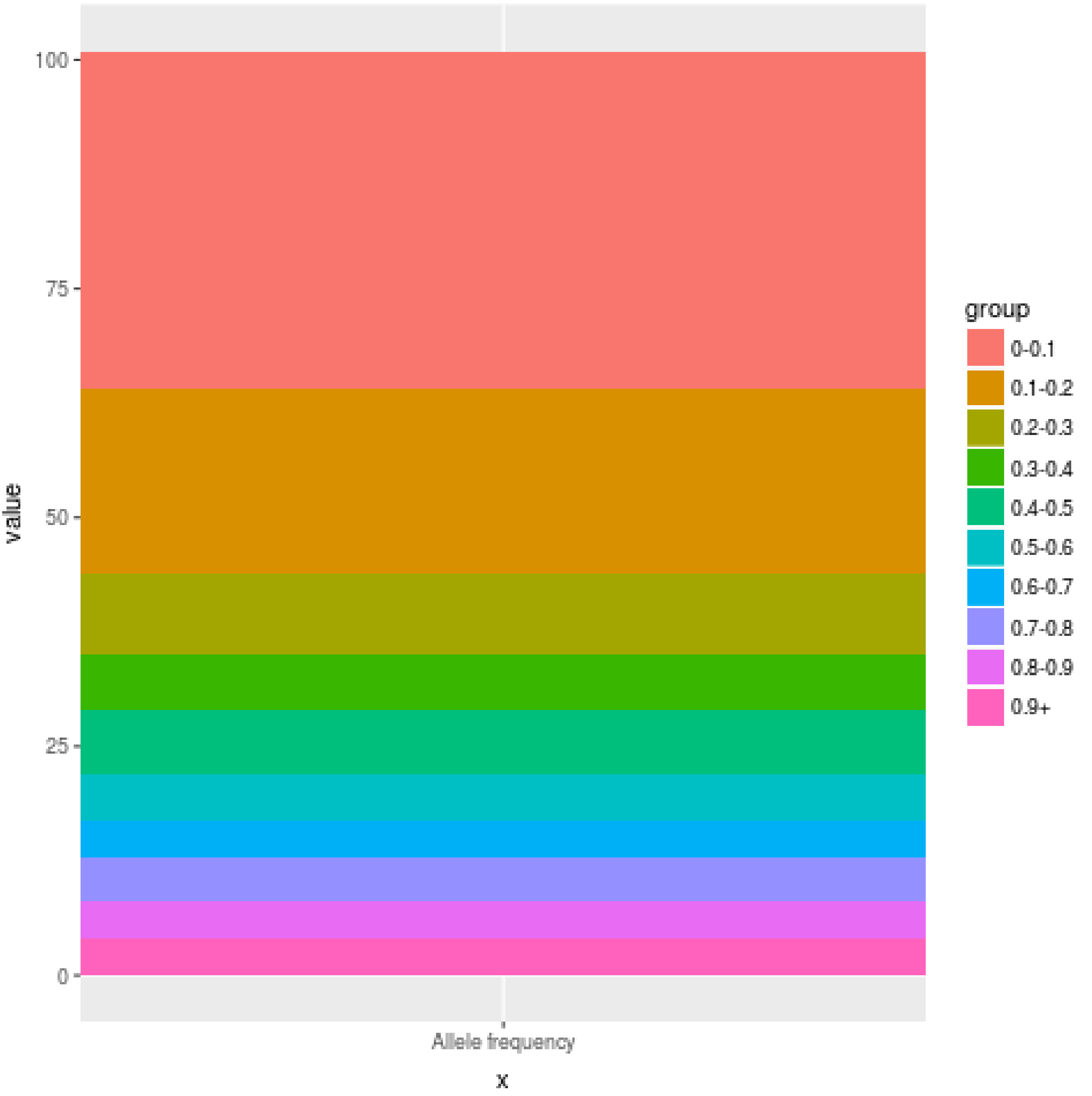

### Supplementary file 4

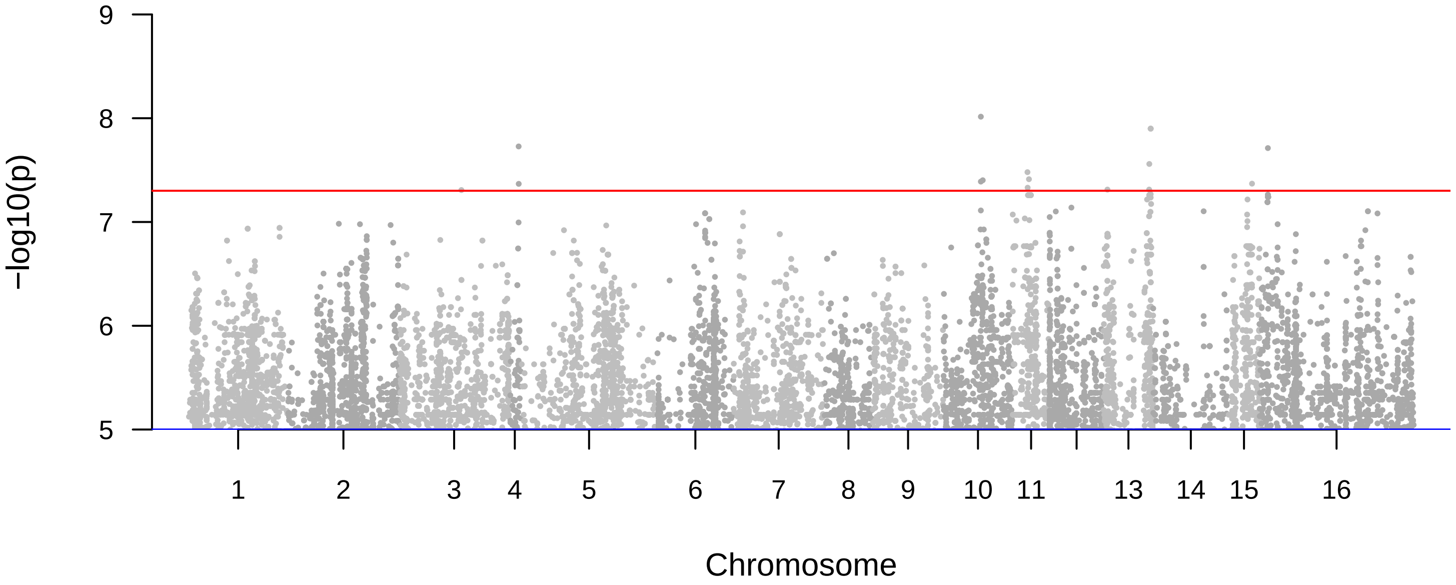

### Supplementary file 5

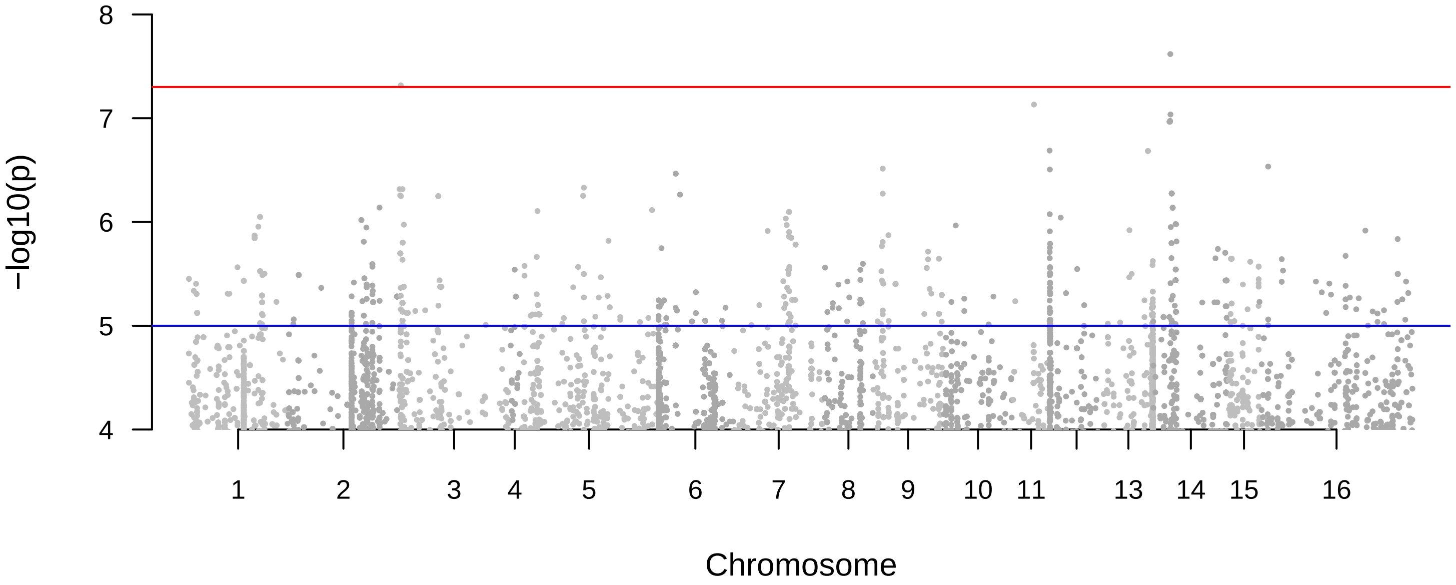

### Supplementary file 6

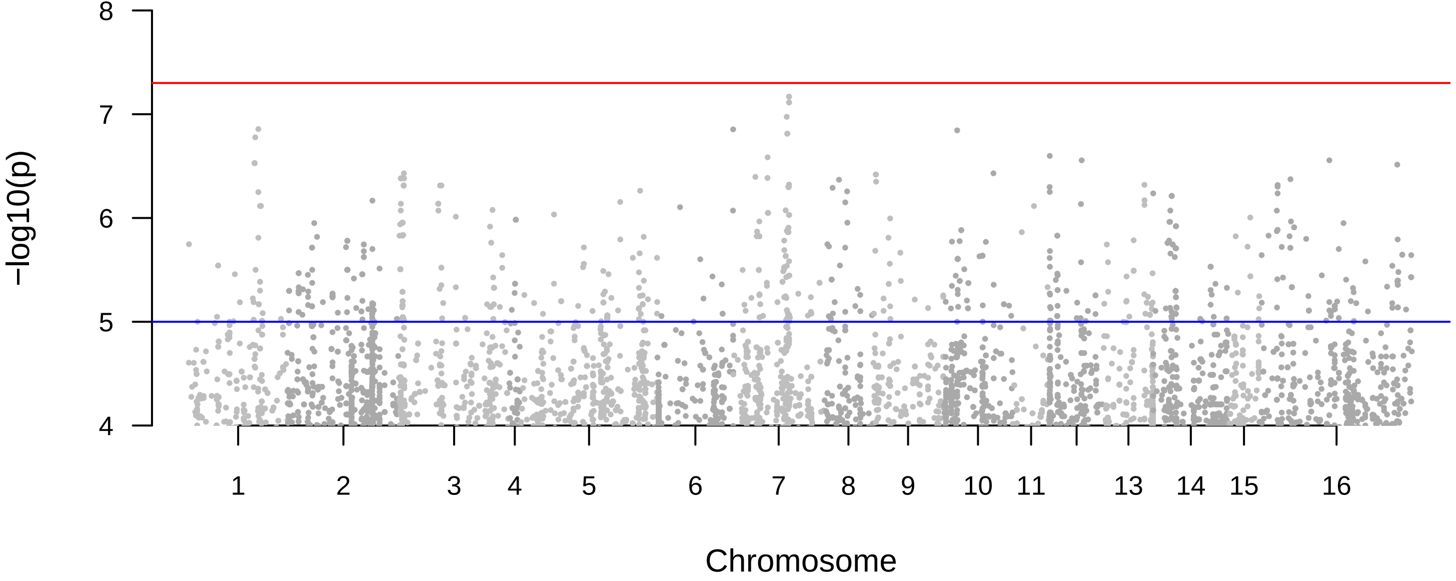
